# Case study of a rhizosphere microbiome assay on a bamboo rhizome with excessive shoots

**DOI:** 10.1101/2021.03.16.435735

**Authors:** Fuqiang Cui, Yifan Yang, Mengyuan Ye, Wei Wei, Wenqian Huang, Ying Wu, Xi Jiao, Xiaoxue Ye, Shutong Zhou, Zhubing Hu, Renyi Gui, Wenwu Wu, Kim Yrjälä, Kirk Overmyer, Shenkui Liu

**Affiliations:** State Key Laboratory of Subtropical Silviculture, Zhejiang A&F University, Lin’an 311300, Hangzhou, China; School of Agricultural Engineering, Jiangsu University, Zhenjiang 212013, China; Organismal and Evolutionary Biology Research Program, Faculty of Biological and Environmental Sciences, and the Viikki Plant Science Centre, University of Helsinki, P.O. Box 65 (Viikinkaari 1), FI-00014 Helsinki, Finland; Key Laboratory of Plant Stress Biology, School of Life Sciences, Henan University, Kaifeng 475004, China

**Keywords:** Bamboo, rhizosphere, microbial diversity, *Burkholderia*, shoot clusters

## Abstract

Young Moso bamboo shoots are a very popular seasonal food. Bamboo is an important source of income for farmers and the value for cultivation has recently been estimated to $30,000 per hectare. A rare and valuable phenomenon has recently appeared where dozens of adjacent buds within a single Moso bamboo rhizome have grown into shoots. Due to its rarity, this phenomenon, which is of practical importance for the production of edible shoots, has not been scientifically studied. We report the occurrence of a rhizome with 18 shoots, of which the microbiome were analyzed, using rhizomes having one or no shoots as controls. The community of prokaryotes, but not fungi, correlated with the shoot numbers. *Burkholderia* was the most abundant genus, which negatively correlated with rhizome shoot number, while *Clostridia* and *Ktedonobacteria* positively correlated with many shoots. Two *Burkholderia* strains were isolated and their plant-growth promoting activity was tested. The isolated *Burkholderia* strains attenuated the growth of bamboo seedlings. Analysis of collected events of enhanced shoot production in China showed no evidence that enhanced shoot development was heritable. Overall, our data provides a firsthand study on excessive shoot development of bamboo.

## Introduction

The young shoots of Moso bamboo (*Phyllostachys edulis*) are a very popular seasonal food. Moso bamboo shoots are rich in fiber and nutrients, making them a desirable health food with increasing demand ^1,2^. The income from bamboo shoot business has reached $30,000 per hectare in advanced cultivation areas ^3^. Techniques to improve shoot production rely mainly on fertilization management ^4–9^. Moso bamboo propagates vegetatively through rhizomes, and each node of a rhizome possesses a single bud. The bud number in one rhizome can reach from tens to over one hundred, depending on the rhizome length and environmental conditions. Most buds remain dormant; less than 5% develop to be shoots ^10^. Given the energy demands of the extremely rapid growth in bamboo shoots (> 1 meter per day at maximum), the germination of too many shoots from one rhizome is detrimental ^11^; energy waste from competition and decay of excess shoots limits its occurrence under natural conditions. For well-cultivated bamboo farms, where shoots are collected at a very young stage, increased shoot number, however, would be economically desirable as it would bring increased income, and have practical labor-saving advantages: Increased numbers of shoots on a rhizome would improve the visibility of small soil mounds pushing up from young shoots, making randomly distributed underground shoots easier for harvest. Thus, promotion of more buds developed to shoots is of both economic and practical importance.

For Moso bamboo, usually one or two neighboring buds develop to be shoots. Four to six neighboring shoots within a rhizome is uncommon. Only very few cases have been reported (no scientific reports available) where more than ten shoots have developed densely along one rhizome and no research has previously been conducted about multiple shoots. Here we term this rare occurrence of multiple shoots as shoot cluster. Studying shoot clusters may offer key information for farmers to cultivate excessive shoots for better income. The very rare occurrence of shoot clusters restricts the chances of effective study on them. It is yet unknown whether shoot cluster is a genetically heritable phenomenon or an occasional result due to environmental factors.

Studies on plant species other than bamboo have found that bud development may involve interactions between hormone and sugar signaling ^9,12–16^. Diverse microbes live in close association with plants and have a significant influence on plant growth, nutrient uptake, and stress tolerance ^17,18^. The correlation between microbes and bud dormancy has not been revealed, while the production of analogues of plant hormones by microbes might give some suggestions ^19^. For example, most rhizosphere microbes produce auxins, which inhibits axillary buds from outgrowth ^20–22^. Strigolactones and cytokinins involved in bud outgrowth, can also be produced by microbes ^23–26^. Gibberellic acid is the major plant hormone that breaks bud dormancy ^27^. The gibberellins, which are analogs of gibberellic acid, are produced by a variety of plant associated microbes ^28–31^. Bamboo is a perennial plant whose rhizomes coexist with an abundant microbes ^32,33^. It is not known what kind of microbes have effects on the shoot development. To answer this question, the composition of microbes related to bamboo shoots should be illustrated first. However, microbiome assay has not been conducted with shoot clusters of bamboo.

We managed to find a case of shoot cluster of bamboo, and thereby obtained the first chance to study this rare phenomenon. The rhizome soil microbiome was analyzed to test the hypothesis that soil microbes may influence shoot cluster formation. Both bacterial and fungal communities were studied in detail to identify possible bacterial or fungal connections to shoot cluster formation. This study may raise awareness of and contribute toward future research into shoot clusters.

## Materials and Methods

### Sample collection and sequencing

Rhizosphere soils of Moso bamboo (*Phyllostachys edulis*) were collected at the root areas (about 30-40 cm in depth) with sterile bags on the 20^th^ of December 2016 (location at 28°34’54.6”N 119°09’59.0”E). Three samples (around 400 g soil per sample) from each type of rhizomes (samples with 18 shoots and no shoots were from distinct parts within the same rhizome) were collected and stored at +4°C after transport to the laboratory. The soil samples were first sieved to exclude roots and coarse gravel, DNA was isolated with the NucleoSpin® Soil kit (Takara, 740780). The isolated DNA was examined by running in 1.0% agarose gel and the concentration was estimated spectrophotometrically (NanoDrop 2000; www.thermofisher.com). For bacteria, the V4 16S rRNA region was amplified using the primers 515F 5’-GTGCCAGCMGCCGCGGTAA-3’ and 806R 5’-GGACTACHVGGGTWTCTAAT-3’ ^34^. For fungi, the rDNA internal transcribed spacer (ITS) region was amplified with primers ITS1F 5’-CTTGGTCATTTAGAGGAAGTAA-3’ and ITS2R 5’-GCTGCGTTCTTCATCGATGC-3’ as previously reported ^35,36^. The amplified PCR products were sequenced by the Tianjin Novogene Bioinformatics Technology Co., Ltd using paired-end sequencing of the Illumina HiSeq platform.

### Sequence data analysis

Multiplexed raw sequencing data were deconvoluted and quality filtered to remove poor quality reads (average quality score < 25, truncated reads < 50 base pairs, ambiguous bases and frame-shift errors) and potential chimeric sequences with QIIME and Mothur ^37,38^. Sequences from each library were clustered into operational taxonomic units (OTUs) with 3% differences using the Uparse program ^39^. A total of 462,263 prokaryotic OTUs and 389,869 fungal OTUs were obtained and classified with Ribosomal Database Project (RDP) Classifier ^40^. The α-diversity indices (Chao, Ace, Shannon) and β-diversity based on both weighted and unweighted Unifrac were calculated and analyzed with the QIIME and Mothur programs. The principle component analysis (PCA) of the samples was performed using QIIME based on the Jaccard distance ^38^. Linear discriminant analysis (LDA) effect size (LEfSe) was calculated online (https://huttenhower.sph.harvard.edu/galaxy/) to determine the biomarkers with LDA = 3 ^41^.

### Isolation of *Burkholderia* species

To facilitate root dissection, bamboo root segments of the less lignified sections (5 cm from the root tips) were collected and washed thoroughly. The roots were treated according to the isolation procedures described in ^42^. The solution containing microbes was evenly spread on half strength (1/2) PDA medium (SLBT9643, Sigma, USA). Microbial colonies were selected and streaked on a new plate. More than 1,000 single colonies were used in growth-promoting assays with plants. For isolates with significant growth-promoting activity 16S rDNA was amplified using primers 27F and 1492R, and sequenced. The sequence data of two isolated *Burkholderia* strains is listed in Supplementary table 4. For the growth-promotion assay, candidate strains were streaked on 1/2 MS medium with rice or bamboo seedlings in aseptic *in vitro* culture without any contact to the seedlings. The seedlings were photographed and weighted after one week with or without microbial strains.

### Growth-promotion assay of rice and bamboo seedlings

To test rice and bamboo seedling *in vitro*, seeds were sterilized with 70% ethanol plus 2% triton X-100 for 3 min, and additionally with 5% sodium hypochlorite for 20 min. The endosperms of sterilized bamboo seeds were excised to eliminate bacterial contamination. Bamboo embryos or intact rice seeds were placed on 1/2 MS medium with 1% sucrose and 1% agar. The isolated strains of *B*. sp. YF and MY were streaked on the medium to avoid direct contact with the sterilized seeds. Using microbe-free medium as control. *B*. sp. YF and *B*. sp. MY did not exhibit visible influence on the germination of bamboo embryos. One-month-old seedlings were photographed and weighted. For soil-grown seedlings, two inoculation methods were used. One was to inoculate bacteria prior to sowing. The seeds were soaked in bacterial suspensions (OD = 0.6) of *B*. sp. YF and *B*. sp. MY, respectively, for 10 min and then sown in an autoclaved soil mixture of (2:1) peat and vermiculite. The second method applied was to inoculate bacteria after sowing. Bamboo seeds were first sown, then watered three times with bacterial suspensions (OD = 0.6) of *B*. sp. YF or *B*. sp. MY. Growth chamber conditions were 150-200 μmol m^-2^ s^-2^, 60% humidity, 12/12 h (light/dark) photoperiod and 23/18°C (day/night). One-month-old seedlings were dried at 70°C for 24 hours before weight measurement.

### Collection of recorded events of rhizomes with more than ten shoots

In Chinese folklore, events involving the development of dozens of shoots within one rhizome are considered as good omens, which are long-cherished and reported in newspapers. This provided a valuable information resource in the absence of scientifically available documents. We collected these reports and summarized key information (time, shoot number, location), see Supplementary table 1. The locations of these rhizomes were manually marked on the map of Zhejiang province and a color key was added using the R program.

### Statistical analysis

Statistical analysis of the dry weight of seedlings was performed with scripts in R (version 3.0.3). Using the nlme package, a linear mixed model with fixed effects for samples, treatments, and their interaction was fitted to the data, plus a random effect for biological repeats. The model contrasts were estimated with the multcomp package, and the estimated P-values were subjected to single-step P-value correction. A logarithm of the data was taken before modeling to improve the model fit. The P-values of the fresh weight of rice were calculated with a two-tailed Student’s *t*-test using equal variance.

## Results

### Location of an 18 shoot rhizome

The shoot cluster was found in a small grove in Suichang (28°34’54.6”N 119°09’59.0”E), a county in southern Zhejiang province. The sampling location was a cultivation terrace on a mountain slope facing to the southeast (Fig. 1A). The rhizome growth area was relatively flat. This terrain is similar to locations of three other events of shoot cluster according to collected news reports (Supplementary Table 1). The rhizome segment containing the18 shoots was 1.3 meters long and was sampled out of a single rhizome of >5 meters in length. The regions adjacent to the shoot cluster had no shoots. Two shoots were destroyed during excavation and 16 attached (Fig. 1B). Microbial community assays were carried out with rhizosphere soils. Soil samples were collected from the rhizosphere of the rhizomes with 18 shoots (E1-E3; Fig. 1C), the regions of the same rhizome up- and down-stream of the18 shoots (N1-N3; Fig. 1C), and three single shoots from other rhizomes in the same grove (S1-S3; Fig. 1C).

**Fig. 1.**
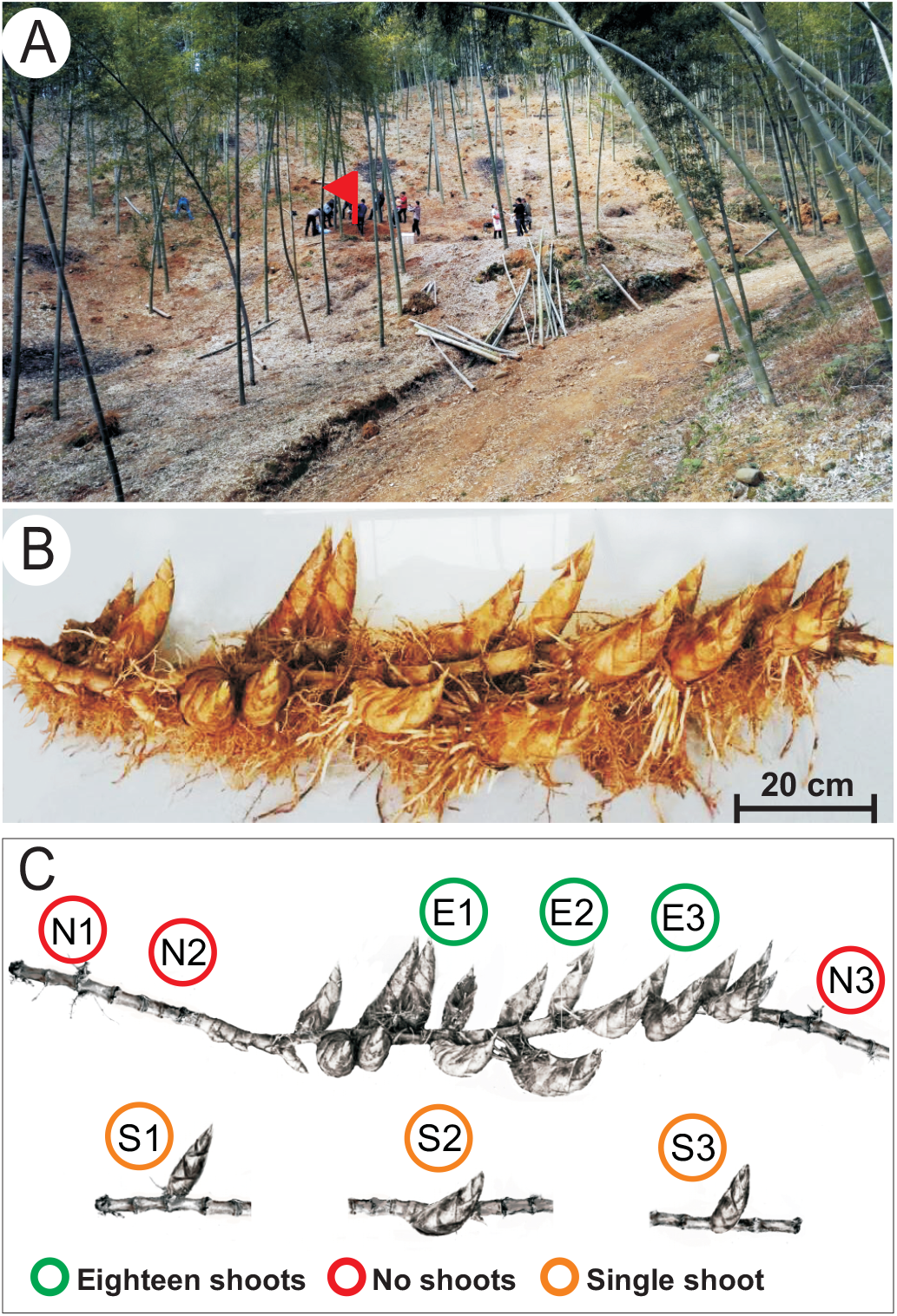
A rhizome with 18 shoots from Moso bamboo plantation was investigated. (A) The terrain of the location. The rhizome growing site is indicated with a red flag. (B) Picture of the rhizome with 18 shoots. Please note that two shoots are missing from the picture as they were destroyed during digging. Bar = 20 cm. (C) Schematic diagram of the positions within the rhizome that were sampled for microbial community analysis. Circles with different colors indicate the sampling positions and types.

### Alpha-diversity of prokaryotic communities

Illumina sequencing of the 16S rRNA amplicon was conducted with DNA isolated and amplified from all rhizosphere soils samples (Fig. 1C). A total 2972 operational taxonomic units (OTUs) were detected with greater than 97% identity to the reference sequences of bacterial 16S rDNA (Supplementary table 2). On average 1694 OTUs were detected from the samples of the region with the shoot cluster, 1536 OTUs from the samples with a single shoot, and 1438 OTUs from the regions with no shoots (Supplementary table 2). This reflected a trend that the higher absolute abundance of prokaryotic OTUs the bigger the number of shoots. This trend was also observed from rarefaction curves of the OTUs (Fig. S1A). In contrast to bacteria, the numbers of fungal OTUs did not show this trend with rhizome shoot number (Supplementary table 3; Fig. S1B). Venn diagrams were constructed with OTUs of the three sample groups with eighteen shoots, a single shoot, and no shoots. The number of prokaryotic OTUs shared between the three samples were 3-10 fold higher than the unique OTUs of each sample (Fig. S2A). A similar trend was also observed between fungal samples; where the shared OTUs were 1.5-6 fold higher than the unique ones (Fig. S2B). This indicated the microbial communities in the rhizosphere were largely stable between samples. The unique prokaryotic OTUs in each sample group exhibited the trend of increasing abundance with increasing shoot numbers (Fig. S2A). The unique fungal OTUs did not exhibit this trend (Fig. S2B).

### Microbial communities of samples with different shoot numbers

Principle component analysis (PCA) demonstrated that prokaryote communities clustered according to sample types, but the fungi did not (Fig. 2A and 2B). The samples with 18 shoots and no shoots were collected from different segments in the same rhizome. The differentiation of prokaryotes was much clearer between these two groups compared to the fungi (Fig. 2A and 2B).

**Fig. 2.**
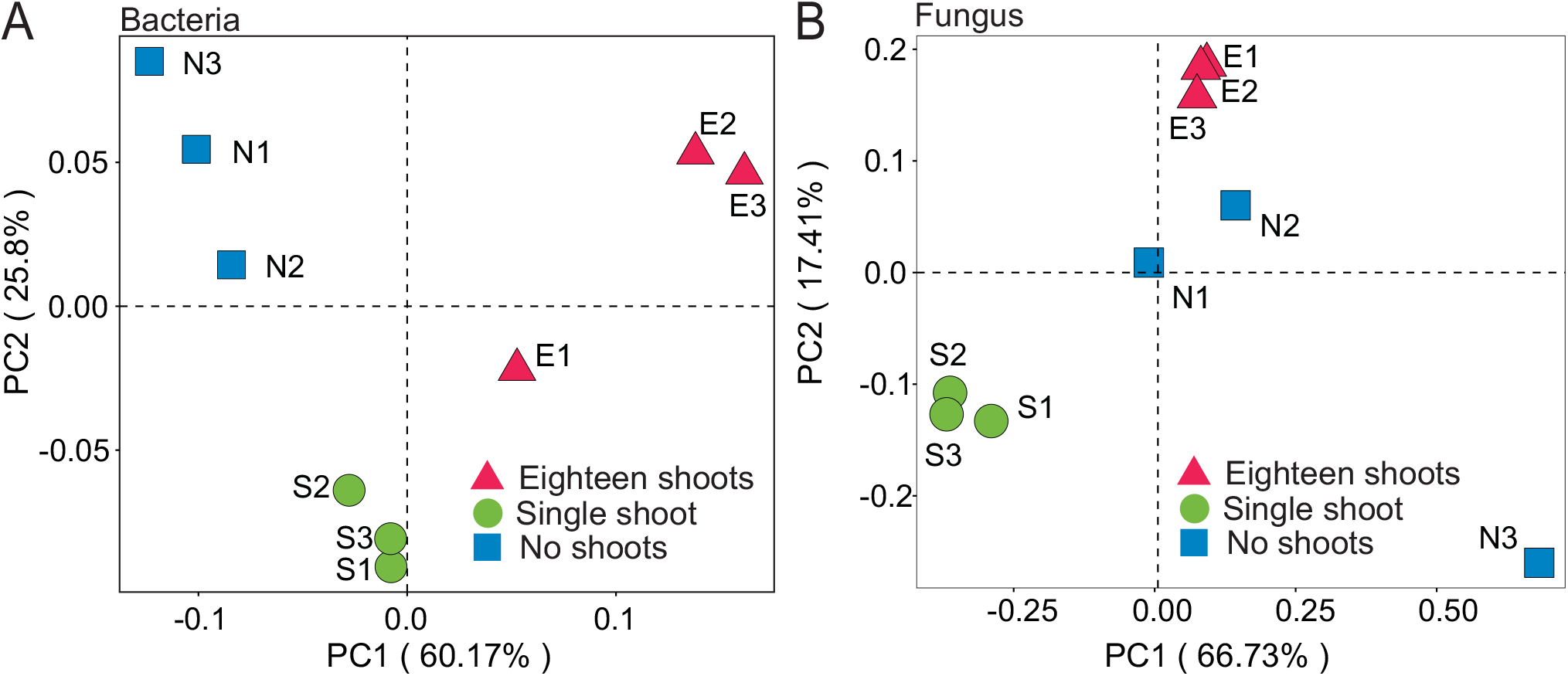
Weighted UniFrac principle component analysis (PCA) of the microbial diversity in three Moso bamboo rhizosphere soil sample types. (A) Prokaryotic 16S rDNA gene sequences. Three repeats of each sample type are indicated with red triangles (18 shoots), green circles (single shoots) and blue square (no shoots). (B) Fungal ITS sequences. Three repeats of each sample type are indicated as in (A).

To identify the predominant discriminant taxa, we performed a linear discriminant analysis (LDA) effect size (LEfSe) analysis ^41^. Representative biomarker taxa were found in each group of samples: classes of *Clostridia* and *Ktedonobacteria* were significantly enriched in samples with many shoots; *Subgroup_3* order was the predominant order of the single shoot samples, while orders *Burkholderiales* and *Legionellales* were predominant orders in samples with no shoots (Fig. 3A). In contrast to prokaryotes, we did not find fungal biomarker taxa for samples with 18 shoots in the LEfSe analysis (Fig. 3B). Only the classes *Eurotiomycetes* and *Agaricomycetes* were significant in samples from no shoots and single shoot, respectively (Fig. 3B). Overall, our data suggest that prokaryotic communities may play a more important role in the regulation of rhizome shoot numbers, while fungal communities were not correlated with shoot numbers. Therefore, we focused on prokaryotic taxa in the following studies.

**Fig. 3.**
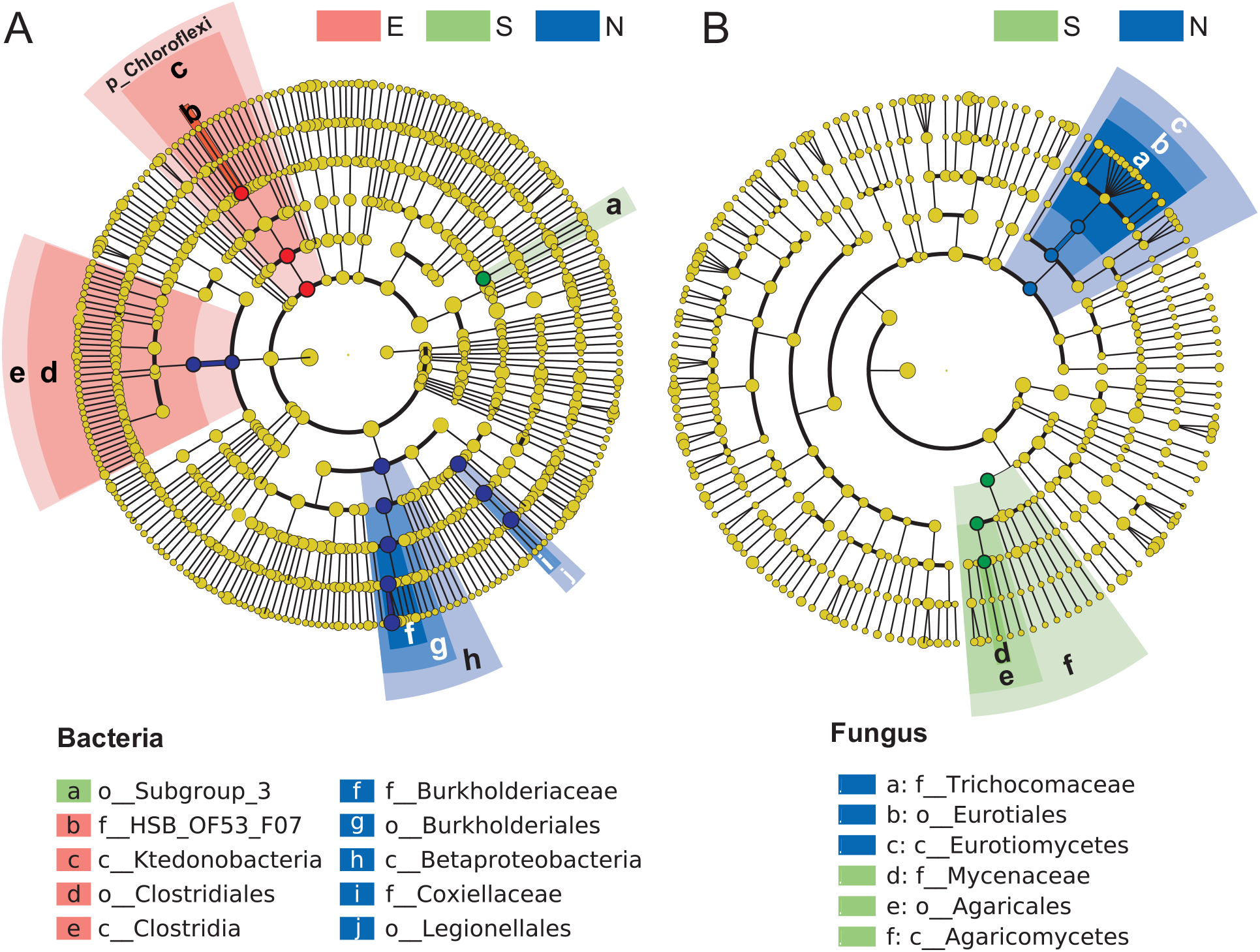
The differential phylogenetic distribution and biomarkers microbial taxa of each group of samples in linear discriminant analysis (LDA) effect size (LEfSe). (A) Bacteria. (B) Fungi. LDA scores ≥ 3. Circles indicate phylogenetic levels from phylum to genus. The yellow nodes represent non-significant differential microbes while the nodes in different colors represent the significant differential biomarker microbes. The color sectors indicate sample types as abbreviated: E, eighteen shoots, in the purple sector; S, single shoot, in the green sector; N, no shoots, in the blue sector.

### The most abundant prokaryotic genera in rhizome soil

The phylogeny of top 100 most abundant genera of bacteria of the data set were further studied by phylogenetic analysis including relative abundance (Fig. 4). The most abundant genus was *Burkholderia* (Fig. 4), which was also the biomarker of samples with no shoots (Fig. 3A). Interestingly, the abundance of *Burkholderia* was negatively correlated with the number shoots in each sample group; the no-shoots sample had the highest abundance, single-shoot was next, and then the 18 shoots (Fig. 4; Fig. S3). The second most abundant genus *Rickettsiella* (Fig. 4) are mainly intercellular arthropod pathogens, which have an opportunistic soil-dwelling habit ^43,44^. Unlike *Burkholderia*, the abundance of *Rickettsiella* exhibited no correlation with the shoot number of each sample group (Fig. 4). Both *Burkholderia* and *Rickettsiella* are proteobacteria, belonging to families of *Burkholderiaceae* and *Coxiellaceae* respectively. These two families were biomarker taxa of samples of no shoots (Fig. 3A). Among the top 100 genera, 22 belonged to the bacterial classes of *Clostridia* and *Ktedonobacteria*, which were prokaryotic biomarker taxa of the 18 shoot samples (Fig. 4; Fig. 3A).

**Fig. 4.**
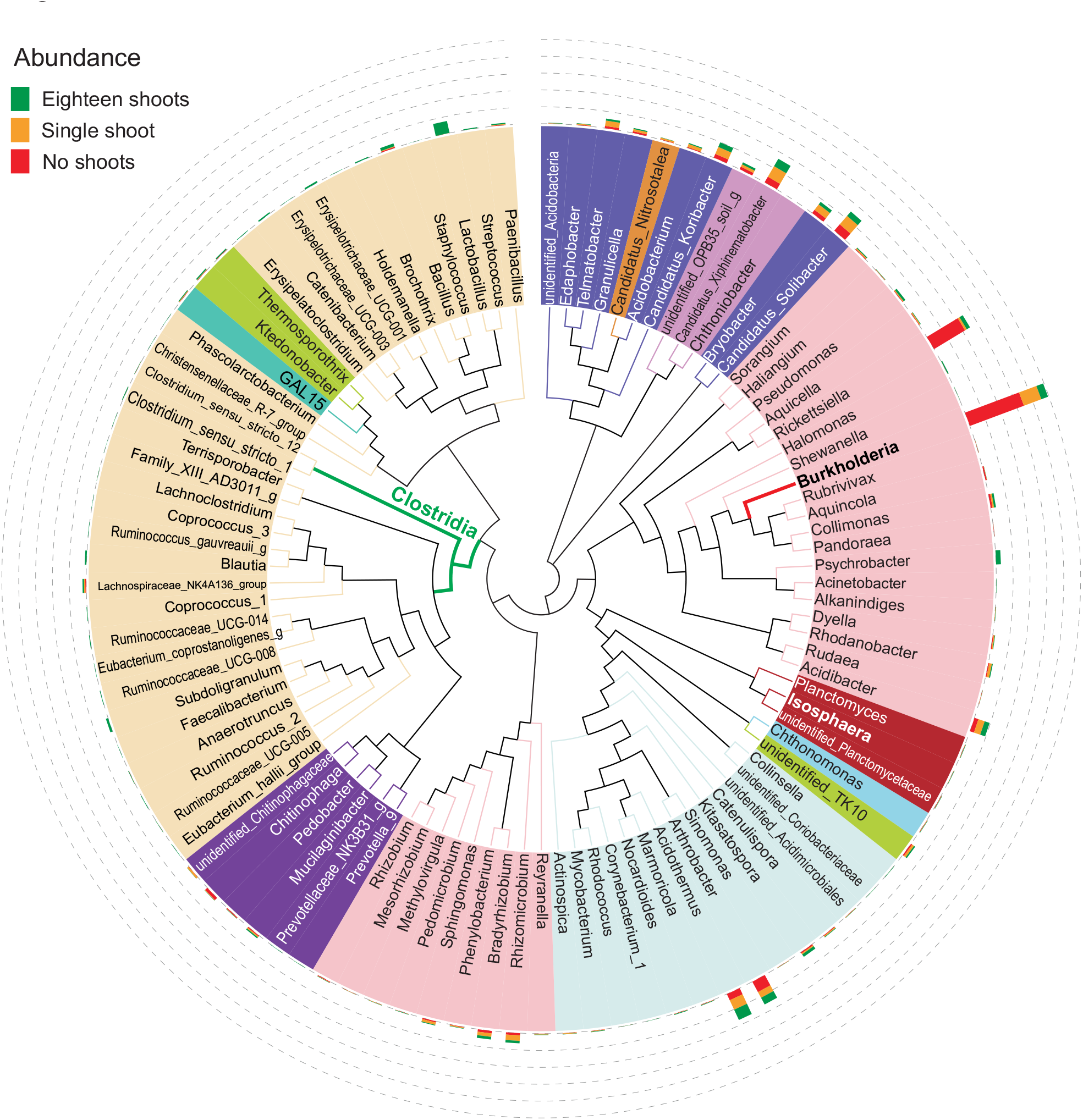
Maximum likelihood phylogeny of the top 100 abundant genera. Phylogenetic tree based on the most abundant 16S sequences was generated using MEGA 5 with the maximum likelihood method. Bootstrap values (500 replicates) of each cluster are given above the branches. Non-coded and coded symbols on tree tips indicate the taxonomic affiliations. The relative abundance of each genus was proportionally represented with the color bar at the outer circles as indicated.

### Isolates of *Burkholderia* species in growth promotion

The isolation of growth promoting microbes from the rhizosphere of our shoot cluster has been conducted as an independent project, which aims to find microbes with potential application of microbial fertilizers for crops, such as rice. The isolation project is beyond the scope of current study, but two *Burkholderia* species were successfully isolated, and were tested here to gain some suggestive insights into the function of *Burkholderia* on bamboo. The two isolates were 99% identical to *Burkholderia* species and further named them *B*. sp. strain YF (*B*. sp. YF) and *B*. sp. strain MY (*B*. sp. MY) respectively (Supplementary table 4). These two *Burkholderia* strains significantly promoted the growth of rice seedlings, as shown visually and quantified in fresh weight (Fig. 5A and 5B). This result is consistent with the well-known ability of *Burkholderia* spp. to promote plant growth ^45^.

**Fig. 5.**
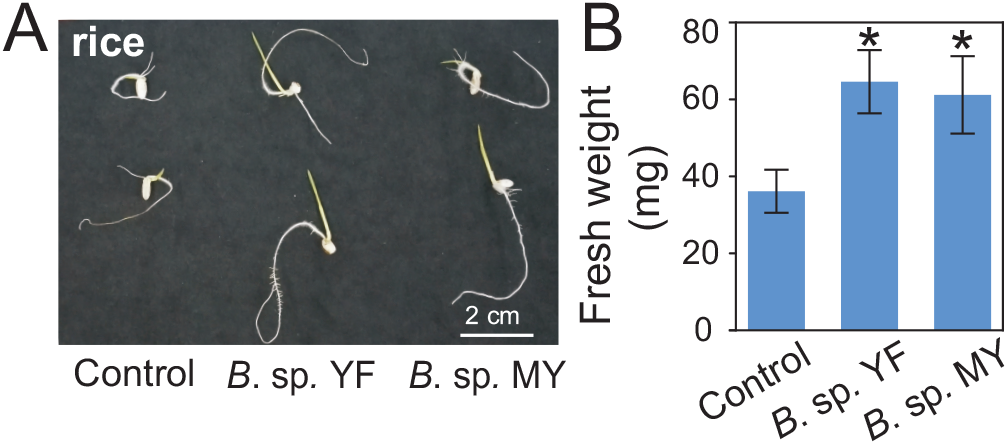
Growth assay of rice seedlings treated with two isolated *Burkholderia* species (*B*. spp.) promote the growth of rice. (A) Rice seedlings grown *in vitro* with or without *B*. sp. YF and *B*. sp. MY. Bar = 2 cm. (B) Quantitative data of fresh weight of rice seedlings treated with *B*. sp. YF and *B*. sp. MY. Errors bars represented SD of the means of 20 replicates. This experiment were repeated twice with similar results. ‘*’ indicates significant groups (*t*-test, *P* < 0.05)

### *Burkholderia* strains tested with bamboo seedlings for growth effects

The growth-promoting activities of *B*. sp. YF and *B*. sp. MY were further tested with *in vitro* grown bamboo seedlings. Neither strain could promote the growth of bamboo seedlings *in vitro* (Fig. 6A and 6B). *B*. sp. YF even attenuated bamboo growth in this assay (Fig. 6A and 6B) compared to control. These results suggested that these two *Burkholderia* strains may not have the ability to promote growth in bamboo. To further confirm this hypothesis, soil grown seedlings were tested. Two inoculation methods were applied; soaking bamboo seeds in bacterial suspensions prior to sowing, or watering seedlings with a bacteria suspensions three times after sowing. Consistent with results from the *in vitro* assay, neither of these two inoculation methods resulted in enhanced growth of bamboo seedlings (Fig. 6C and D). In addition, watering with *B*. sp. YF suspension and pre-soaking seeds with *B*. sp. MY significantly attenuated bamboo growth (Fig. 6C and 6D). Overall, the two isolated *Burkholderia* stains had negative effects on the growth of bamboo seedlings.

**Fig. 6.**
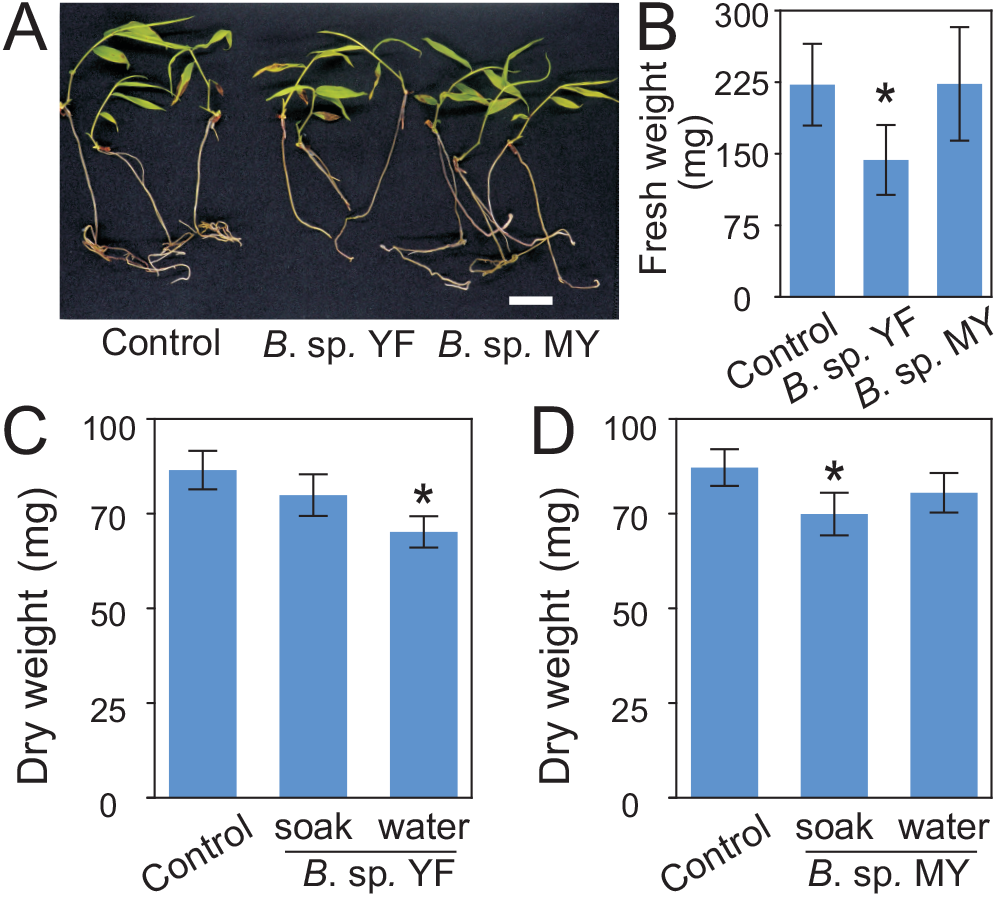
Growth assay of Moso bamboo seedlings treated with *Burkholderia* species *B*. sp. YF and MY. (A) Photographs of one-month-old seedlings with or without application of *B*. sp. YF and *B*. sp. MY. Bar = 2 cm. (B) Quantitative data of fresh weight of the bamboo seedlings in (A). ‘*’ indicates significant groups (*t*-test, *P* < 0.05). (C, D) Soil-grown bamboo seedlings inoculated with or without *B*. sp. YF and *B*. sp. MY. Inoculation was performed with two methods; soaking bamboo seeds in suspension of *B*. sp. YF and *B*. sp. MY respectively before sowing (abbreviated as soak), or watering bamboo seedlings with the bacteria suspension three times during growth (abbreviated as water). Data of three repeats were analyzed in a linear mixed model with single-step P-value adjustment. Error bars represent SE of means. ‘*’ indicates significant groups (*P* < 0.05).

### Collection of reported events of shoot cluster in China

In order to search for clues related to the causes of this rare phenomenon, a total of 18 reported occurrences of shoot clusters were collected, which have been reported in Chinese news media. Events of shoot clusters are considered as good omens in Chinese folklore, which are popular reports in newspapers. Due to the rarity of shoot clusters and no other documents available, newspapers were the only information resource. The collected events are from the period of 2015 to 2019 in all of China (Supplementary table 1). With the exception of four events in two other provinces, all of the events (14 total) were from the Zhejiang province (Supplementary table 1), one of the most advanced regions for bamboo cultivation ^46^. The locations of these 14 events are indicated on the map of Zhejiang province (Fig. 7A). The numbers of shoots per rhizome and the years of each event were also indicated with colors and different shapes (Fig. 7A). These events were randomly distributed over the entire province with no events occurring twice at the same location (Fig. 7A). This suggested that the phenomenon of shoot cluster takes place opportunistically and unrepeatably. This supports the idea that formation of shoot clusters is not genetically heritable. The timing of events varied: six times in 2016, five in 2017, three in 2015, and only twice each in 2018 and 2019 (Fig. 7B). The average shoot number over all of the years was similar (Fig. 7B).

**Fig. 7.**
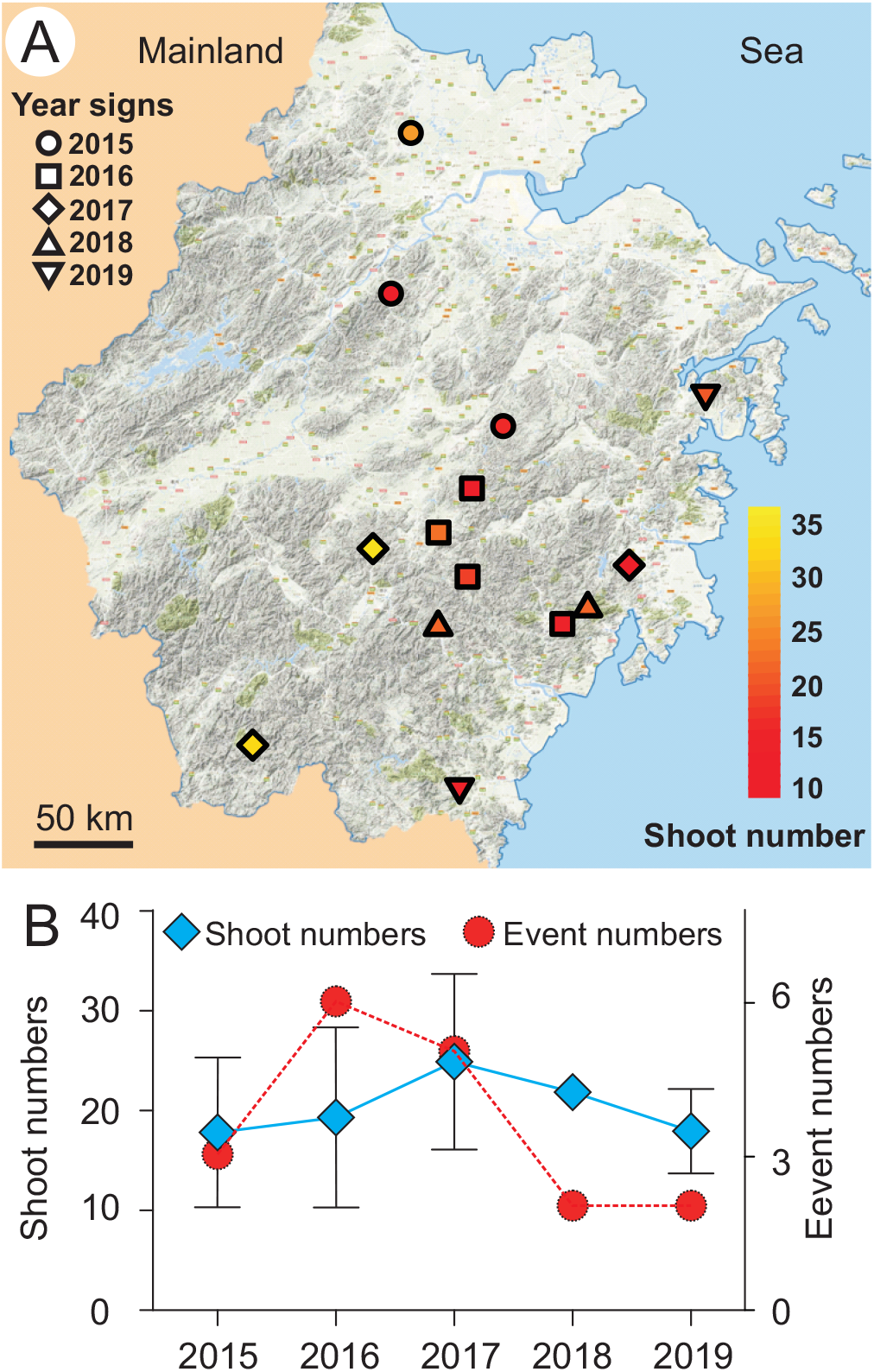
Collection of events of rhizomes with dozens of shoots in Zhejiang province. A total 14 events were recorded. (A) The location of the collected events were marked on the map of Zhejiang province in the East China. The recorded events were from the years 2015-2019, which are represented with different shapes as indicated. The shoot number of each rhizome was from 11-36 as indicated with the color key from red to yellow. Scale bar = 50 km. (B) The number of events recorded each year and the average shoot number of the events for each year were presented. Red circle indicates event number. Blue rhombus indicates average shoot number of each year.

## Discussion

Bacterial and fungal community structures in the bamboo rhizosphere soils were studied here. Principle component analysis (PCA) demonstrated that prokaryote communities clustered according to sample types, while the fungi did not. Closer study of bacterial communities showed that classes of *Clostridia* and *Ktedonobacteria* were significantly enriched in samples with many shoots. These results suggest that bacteria somehow interact with the bamboo in shoot formation. The bacteria *Burkholderia* was predominant in no shoot samples, and correlated negatively with shoot numbers. All these data were solid first-hand information for studying shoot cluster, a agriculturally valuable phenomenon. We discuss the possible correlations between our data and shoot cluster below, which might provide perspective insights for further studies.

Bamboo shoots have a special importance in Southern China where fresh shoots are a desired seasonal food and that generates much needed income for many farmers especially in rural areas. Multiple shoots are rarely formed from one main root (rhizome). We were fortunate to obtain the opportunity to investigate this rare event utilizing microbial ecology tools, with the aim to scientifically document this phenomenon and collect primary information as the basis for hypothesis testing in future studies. The total lack of reports about shoot clusters could be explained by their rarity, but also unpredictability. They are usually discovered by local farmers who lack awareness or interest in the scientific value of the phenomena, not giving researchers notice of the opportunity for further study. Shoot cluster events seem also to have appeared increasingly in recent years according to the literature records (Supplementary table 1). The extreme cases of shoot clusters collected here may facilitate investigation into typical conditions that are required for the induction of shoot development.

Bacteria of the classes *Clostridia* and *Ktedonobacteria* were the marker prokaryotes in the microbial communities associated with clustered shoots (Fig. 3A). Among the top 100 most abundant genera, 22 belonged to these two classes (Fig. 4). Species in the genus *Clostridium*, belonging to *Clostridia*, have been reported to produce gibberellins, which are essential phytohormones for bud germination ^27,47^. Members of *Clostridium sensu stricto* are the true representatives cluster of *Clostridium* ^48–50^. *Clostridium sensu stricto* 1 and 12 were among the top 100 most abundant genera (Fig. 4). *Ktedonobacteria* is a newly established class with relatively large genomes, complex metabolism, and aerobic lifestyles ^51^. Studies on this class are often related to their antibiotic production ^52,53^. Some bacteria in this class, such as *Streptomyces*, exhibited plant growth promoting activity ^54^. The *Ktedonobacteria*, have not yet been tested for plant-growth modulating activity. Recently, *Ktedonobacterial* strains were isolated from decayed bamboo stems^52,55^, indicating *Ktedonobacteria* genera might influence bamboo via a way rather than growth-promotion. The effect of bacteria on plant development are usually a combined result of several different bacterial species. The elucidation of the function of *Clostridia* and *Ktedonobacteria* in bamboo bud germination, will require significant efforts for bacteria isolation and bamboo physiological assays, which is beyond the scope of the current study.

*Burkholderia* was negatively correlated with the number shoots in each sample group. Many *Burkholderia* species have plant growth-promoting activities ^56,57^. Our two isolated *Burkholderia* strains from bamboo rhizosphere were also growth promoting in rice (Fig. 5). However, co-cultivation with these *Burkholderia* attenuated the growth of bamboo seedlings (Fig. 6). This negative effects of *Burkholderia* on bamboo growth is consistent with our result that the abundance of *Burkholderia* were negatively correlated to shoot numbers of rhizomes (Fig. 4 and Fig. S3), suggesting a possible function pattern of *Burkholderia* on shoot formation. It remains unclear why these two *Burkholderia* strains could promote the growth of rice, but not bamboo. One interesting observation was that rice seedlings developed increased root hairs upon *Burkholderia* treatment while bamboo seedling did not (Fig. 6 and unpublished data). Root hairs are essential for establishment of beneficial association between plant and rhizosphere bacteria ^58,59^. Lack of root hairs may prevent bamboo seedlings from receiving benefits from the *Burkholderia* stains. There are also other possibilities for the negative regulation of *Burkholderia* on bamboo shoot numbers. For example, ethylene is required for bud dormancy release in several plants ^60–62^. *Bulrkholderia* species produce 1-aminocyclopropane-1-carboxylate (ACC) deaminase that decrease the ethylene level in plants ^57^. The hypothesis would then be that deaminase production could prevent dormancy release in bamboo, which will require experimental support.

Many environmental factors may be also involved in the regulation of bamboo bud germination, which were collected here in order to provide a whole information of shoot cluster study so far. The terrain where the shoot cluster set is relatively flat on a slope. Flat areas on slopes allow deposition of nutrients brought from soils above by rainwater. The retention of rainwater flow in the flat areas may also result in hypoxic conditions, which would benefit the propagation of anaerobic bacteria. *Clostridia*, the marker genera of shoot cluster in this study are obligate anaerobes. Temperature might also be a factor. The occurrence of shoot clusters between years varies largely in our collection. It peaks in 2016 with six events (Fig. 7B). January 2016 was the coldest winter in recent 30 years in China, during which numerous plants were killed by the fierce freeze ^63^. Freezing exposure has been reported to release bud dormancy in multiple perennial trees ^64–66^. It would make sense that the short exposure to freezing temperatures in 2016 could have promoted shoot cluster occurrence. Environmental factors would however influence areas on a large scale, while reported shoot cluster events have remained sporadic and isolated. The opportunistic character of microbes might be more suitable to explain the random and rare occurrence of shoot clusters.

Overall, this is the first study on the rare phenomenon of shoot clusters. We analyzed the microbial community, recorded the terrain of the locations where it occurs, and discussed the influence of environmental factors. This information provides a reference point for future studies of this previously understudied topic.

## Acknowledgements

This work supported by the Qianjiang Talent D program to FC; the National Natural Science Foundation of China (grant no. 31700224; 31871233; 31770543); the Zhejiang Science and Technology Major Program on Agricultural New Variety Breeding (grant no. 2016C02056-1); the Program for Changjiang Scholars and Innovative Research Team in University (grant no. IRT_17R99 to SL). KO is funded by the Academy of Finland Center of Excellence in Primary Producers 2014-2019 (decisions #271832 and 307335).

## Author Contributions

FC designed the experiments and conceived the data. WeiWei and XY performed analysis of the sequence data. YY, MY, WH, YW and SZ isolated the *Burkholderia* species and performed plant treatments. SL, RG, WW, FC collected the soil samples. XJ prepare the samples for sequencing. FC, KO, KY and ZH wrote the manuscript with all authors’ edits and approval.

## Legends

**Fig. 1. A rhizome with 18 shoots from Moso bamboo plantation was investigated**. (A) The terrain of the location. The rhizome growing site is indicated with a red flag. (B) Picture of the rhizome with 18 shoots. Please note that two shoots are missing from the picture as they were destroyed during digging. Bar = 20 cm. (C) Schematic diagram of the positions within the rhizome that were sampled for microbial community analysis. Circles with different colors indicate the sampling positions and types.

**Fig. 2. Weighted UniFrac principle component analysis (PCA) of the microbial diversity in three Moso bamboo rhizosphere soil sample types**. (A) Prokaryotic 16S rDNA gene sequences. Three repeats of each sample type are indicated with red triangles (18 shoots), green circles (single shoots) and blue square (no shoots). (B) Fungal ITS sequences. Three repeats of each sample type are indicated as in (A).

**Fig. 3. The differential phylogenetic distribution and biomarkers microbial taxa of each group of samples in linear discriminant analysis (LDA) effect size (LEfSe)**. (A) Bacteria. (B) Fungi. LDA scores ≥ 3. Circles indicate phylogenetic levels from phylum to genus. The yellow nodes represent non-significant differential microbes while the nodes in different colors represent the significant differential biomarker microbes. The color sectors indicate sample types as abbreviated: E, eighteen shoots, in the purple sector; S, single shoot, in the green sector; N, no shoots, in the blue sector.

**Fig. 4. Maximum likelihood phylogeny of the top 100 abundant genera**. Phylogenetic tree based on the most abundant 16S sequences was generated using MEGA 5 with the maximum likelihood method ^67^. Bootstrap values (500 replicates) of each cluster are given above the branches. Non-coded and coded symbols on tree tips indicate the taxonomic affiliations. The relative abundance of each genus was proportionally represented with the color bar at the outer circles as indicated.

**Fig. 5. Growth assay of rice seedlings treated with two isolated *Burkholderia* species (*B*. spp.)**. (A) Rice seedlings grown *in vitro* with or without *B*. sp. YF and *B*. sp. MY. Bar = 2 cm. (B) Quantitative data of fresh weight of rice seedlings treated with *B*. sp. YF and *B*. sp. MY. Error bars represented SD of the means of 20 replicates. This experiment were repeated twice with similar results. ‘*’ indicates significant groups (*t*-test, *P* < 0.05)

**Fig. 6. Growth assay of Moso bamboo seedlings treated with *Burkholderia* species *B*. sp. YF and MY**. (A) Photographs of one-month-old seedlings with or without application of *B*. sp. YF and *B*. sp. MY. Bar = 2 cm. (B) Quantitative data of fresh weight of the bamboo seedlings in (A). ‘*’ indicates significant groups (*t*-test, *P* < 0.05). (C, D) Soil-grown bamboo seedlings inoculated with or without *B*. sp. *YF* and *B*. sp. *MY*. Inoculation was performed with two methods; soaking bamboo seeds in suspension of *B*. sp. YF and *B*. sp. MY respectively before sowing (abbreviated as soak), or watering bamboo seedlings with the bacteria suspension three times during growth (abbreviated as water). Data of three repeats were analyzed in a linear mixed model with single-step P-value adjustment. Error bars represent SE of means. ‘*’ indicates significant groups (*P* < 0.05).

**Fig. 7. Collection of events of rhizomes with dozens of shoots in Zhejiang province**. A total 14 events were recorded. (A) The location of the collected events were marked on the map of Zhejiang province in East China. The recorded events were from the years 2015-2019, which are represented with different shapes as indicated. The shoot number of each rhizome was from 11-36 as indicated with the color key from red to yellow. Scale bar = 50 km. (B) The number of events recorded each year and the average shoot number of the events for each year were presented. Red circle indicates event number. Blue rhombus indicates average shoot number of each year.

**Supplementary Fig. 1. Rarefaction curves of OTUs at 97% similarity for samples from eighteen Moso bamboo shoots, single shoot, and no shoots**.

(A) Bacteria. (B) Fungi. Samples from eighteen shoots, single shoot, and no shoots are indicated by red triangle, green circle, and blue square respectively. The average OTUs number of three samples of each group are shown.

**Supplementary Fig. 2. Venn diagram of OTUs of each group of rhizosphere soil samples**. (A) Bacteria. (B) Fungi. Abbreviations: N, no shoots; S, single shoots; E, eighteen shoots.

**Supplementary Fig. 3. Relative abundance of *Burkholderia* bacteria in each replicate**. The abundance in each replicate of the most abundant genera *Burkholderia* in Fig. 4 are shown individually. The bar in each group represents the average abundance of the three samples. Abbreviations: N, no shoots; S, single shoots; E, eighteen shoots.

**Supplementary table 1. Collection of events of shoot clusters in China.**

**Supplementary table 2. OTUs of bacteria from each sample listed in Fig. 1C.**

**Supplementary table 3. OTUs of fungi from each sample listed in Fig. 1C**.

**Supplementary table 4. The 16S rDNA sequence of the isolated *Burkholderia* species**.

**Supplementary Fig. 1.**
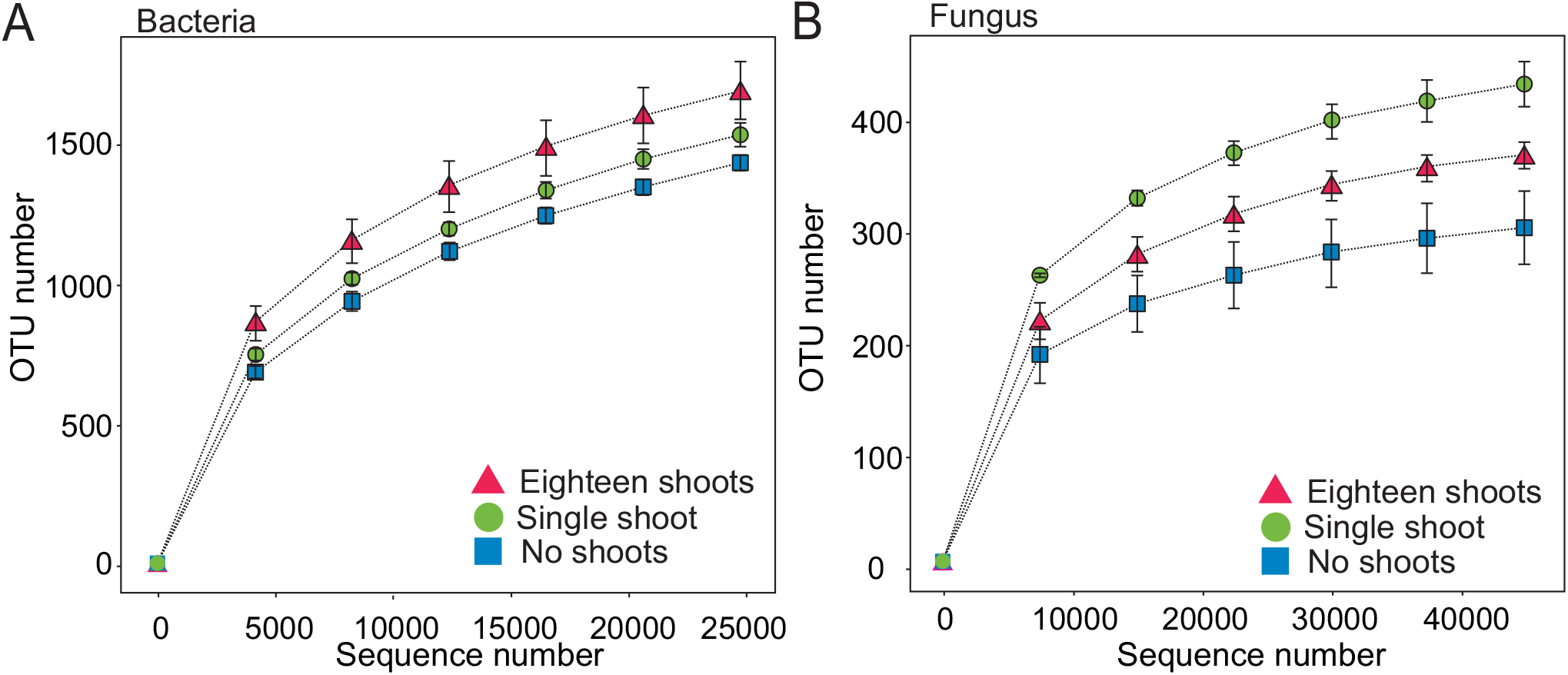
Rarefaction curves of OTUs at 97% similarity for samples from eighteen Moso bamboo shoots, single shoot, and no shoots. (A) Bacteria. (B) Fungi. Samples from eighteen shoots, single shoot, and no shoots are indicated by red triangle, green circle, and blue square respectively. The average OTUs number of three samples of each group are shown.

**Supplementary Fig. 2.**
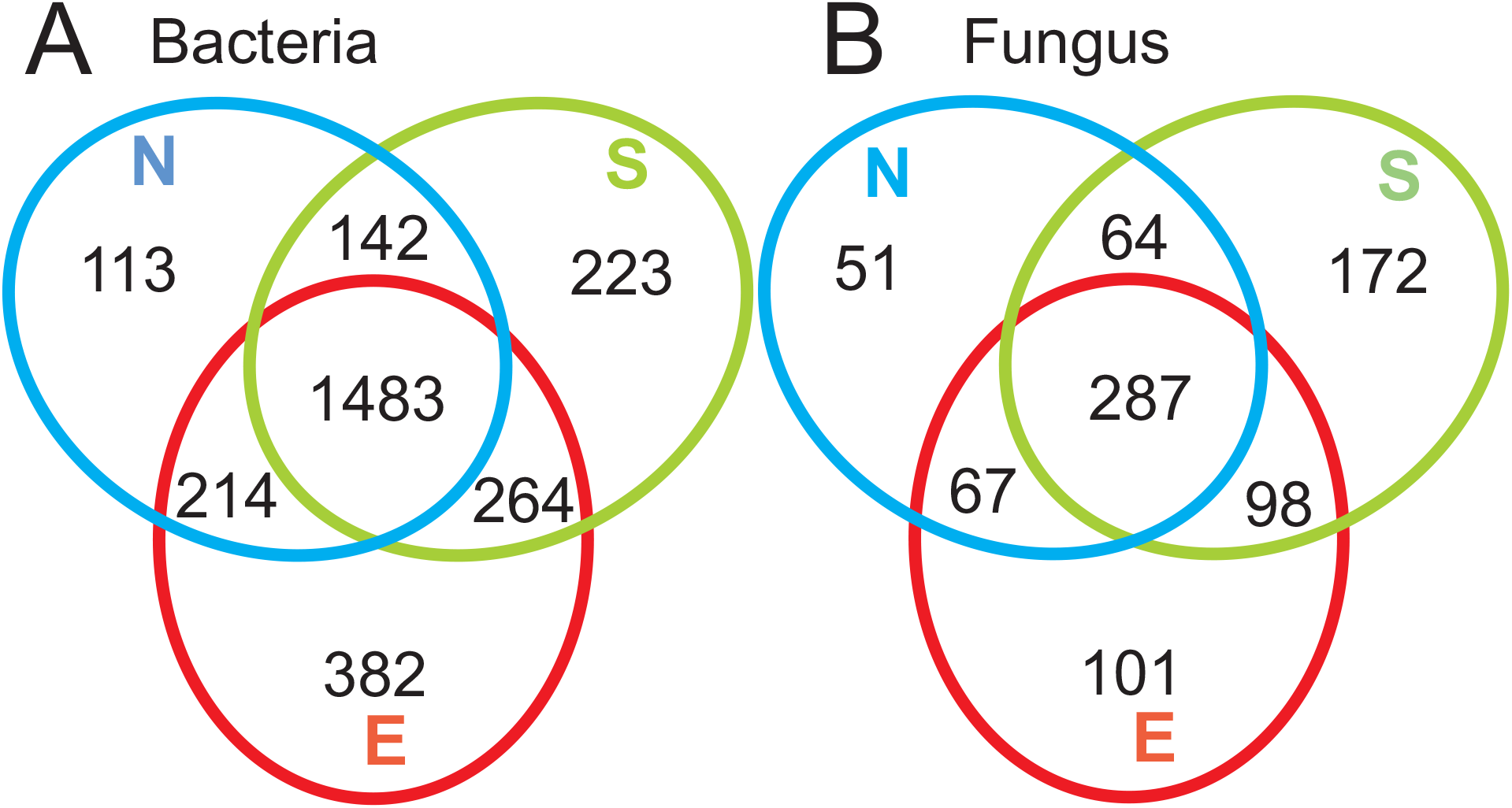
Venn diagram of OTUs of each group of rhizosphere soil samples. (A) Bacteria. (B) Fungi. Abbreviations: N, no shoots; S, single shoots; E, eighteen shoots.

**Supplementary Fig. 3.**
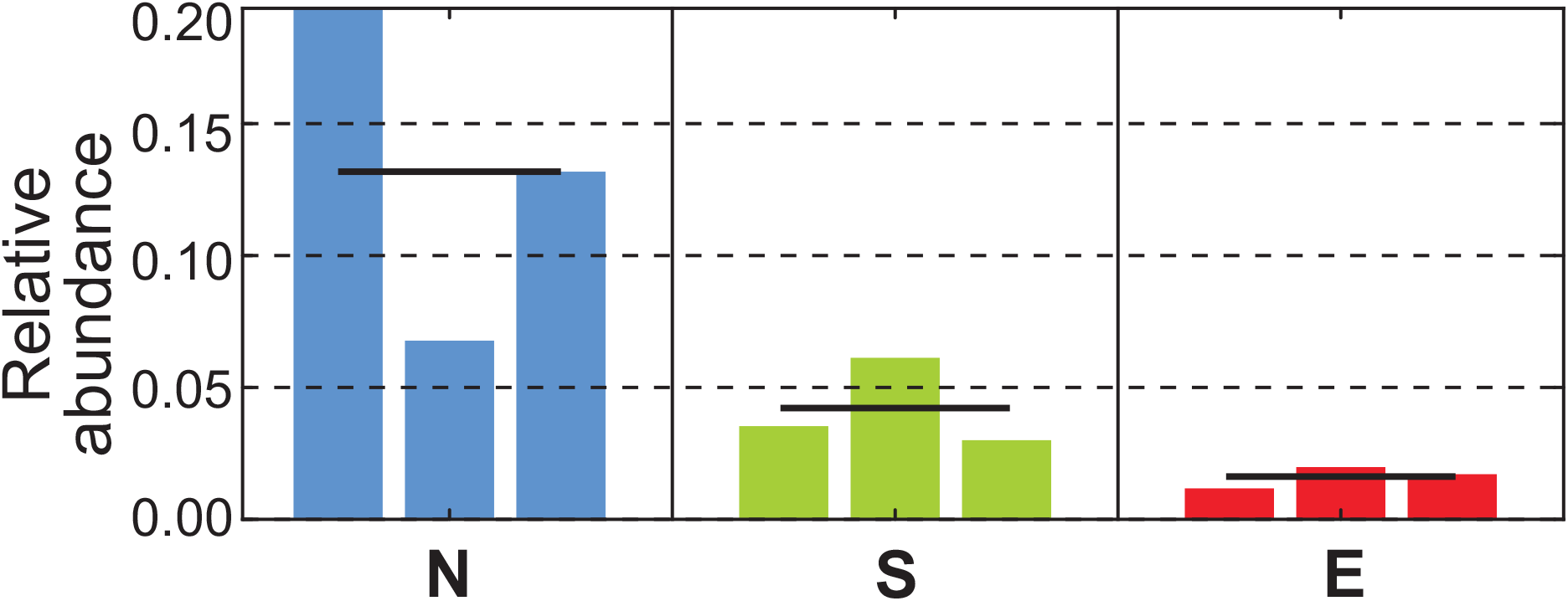
Relative abundance of *Burkholderia* bacteria in each replicate. The abundance in each replicate of the most abundant genera *Burkholderia* in Fig. 4 are shown individually. The bar in each group represents the average abundance of the three samples. Abbreviations: N, no shoots; S, single shoots; E, eighteen shoots.

